# Structural templates for imaging EEG cortical sources in infants

**DOI:** 10.1101/2020.06.20.162131

**Authors:** Christian O’Reilly, Eric Larson, John E. Richards, Mayada Elsabbagh

**Author notes:** Corresponding author: Christian O’Reilly, Ph.D., 3775 Rue University, Room C18, Duff Medical Building, Montréal, Québec, H3A 2B4, 1-438-933-3696. Contributions: COR: Conceptualization; Methodology; Software; Writing - Original draft, review & editing EL: Methodology; Software; Writing - Review & Editing JR: Methodology; Resources; Writing - Review & Editing ME: Resources; Funding acquisition; Writing - Review & Editing.

## Abstract

Electroencephalographic (EEG) source reconstruction is a powerful approach that helps to unmix scalp signals, mitigates volume conduction issues, and allows anatomical localization of brain activity. Algorithms used to estimate cortical sources require an anatomical model of the head and the brain, generally reconstructed using magnetic resonance imaging (MRI). When such scans are unavailable, a population average can be used for adults, but no average surface template is available for cortical source imaging in infants. To address this issue, this paper introduces a new series of 12 anatomical models for subjects between zero and 24 months of age. These templates are built from MRI averages and volumetric boundary element method segmentation of head tissues available as part of the Neurodevelopmental MRI Database. Surfaces separating the pia mater, the gray matter, and the white matter were estimated using the Infant FreeSurfer pipeline. The surface of the skin as well as the outer and inner skull surfaces were extracted using a cube marching algorithm followed by Laplacian smoothing and mesh decimation. We post-processed these meshes to correct topological errors and ensure watertight meshes. The use of these templates for source reconstruction is demonstrated and validated using 100 high-density EEG recordings in 7-month-old infants. Hopefully, these templates will support future studies based on EEG source reconstruction and functional connectivity in healthy infants as well as in clinical pediatric populations. Particularly, they should make EEG-based neuroimaging more feasible in longitudinal neurodevelopmental studies where it may not be possible to scan infants at multiple time points.

**Graphical abstract:** 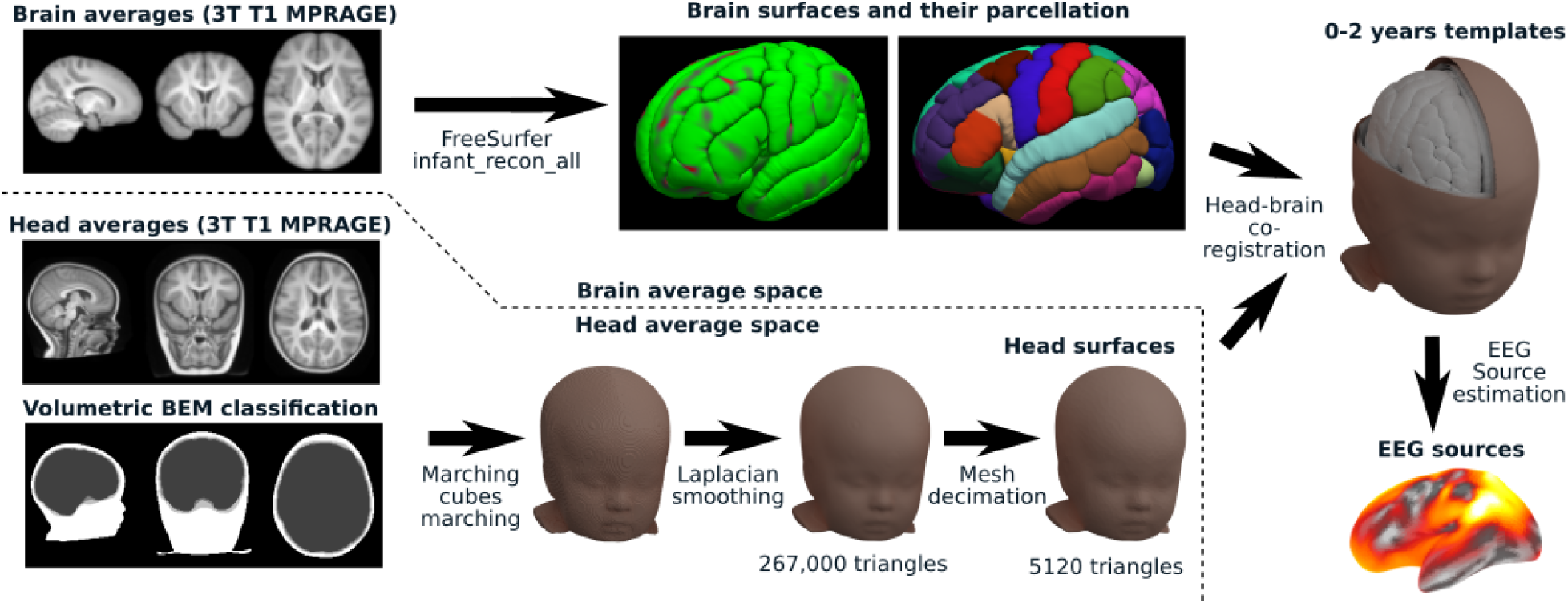

**Highlights:**

- Twelve surface templates for infants in the 0-2 years old range are proposed
- These templates can be used for EEG source reconstruction using existing toolboxes
- A relatively modest impact of age differences was found in this age range
- Correlation analysis confirms increasing source differences with age differences
- Sources reconstructed with infants versus adult templates significantly differ

## Introduction

Our ability to study the functional connectivity of the brain during its early years is crucial in understanding both neurotypical and abnormal developmental trajectories, such as in the case of attention deficit hyperactivity disorder (Konrad and Eickhoff, 2010) or autism (O’Reilly et al., 2017). However, most EEG studies in infants are performed at the scalp level, which presents severe limitations such as the obfuscated relationship between the scalp location of EEG activity and the neuronal sources (Nunez and Srinivasan, 2006; Van de Steen et al., 2019), the impact of the recording reference (Bringas Vega et al., 2019; Guevara et al., 2005), and the confounding effect of volume conduction on EEG power and functional connectivity (Nunez et al., 1997; O’Reilly and Elsabbagh, 2020; Van de Steen et al., 2019). Many of these limitations can be mitigated by unmixing the scalp activity and estimating its neuronal sources.

EEG sources can be estimated in surfaces or in volumes, depending on how dipolar sources are allowed to be placed in the model of the head. The reconstruction of cortical sources using surfaces relies on our understanding of the generative mechanism of EEG to constrain the position and the orientation of dipolar sources. The proximity of the cortex to the scalp, the creation of a dipolar source between the apical tree and the soma of pyramidal cells following postsynaptic depolarization of apical dendrites, and the creation of an open field by the parallel alignment of the apical dendrites are a few reasons that led researchers to hypothesize that scalp EEG is mainly reflecting highly synchronous postsynaptic currents generated in apical trees of cortical pyramidal cells (Baillet et al., 2001). Accordingly, it is a common practice to estimate EEG cortical sources by fitting the amplitude of dipoles whose position and orientation are constrained by the cortical surface.

Alternatively, all dipolar contributions can be postulated to sum linearly in equivalent current dipoles (ECD), which position and orientation is generally left free. Such an approach has some analytic advantages (e.g., mathematical tractability when used with spherical head models) and can be motivated when one or a few specific sources of activity clearly dominates like in the context of epileptic activity (Ebersole, 1994) or for localized independent components (Acar et al., 2016). By generalizing the ECD approaches to a large number of dipoles and using sophisticated finite element models (FEM) of the head, we can also perform current density reconstruction (CDR). These volumetric approaches of source estimation position dipoles using less a priori constraints than methods using cortical surfaces (e.g., dipoles position may not be limited to grey matter or cortical regions, dipole orientation may be free).

Surface and volume source estimation can also be combined, for example by using physiological hypotheses to constrain dipoles within the grey matter of the cortex to be aligned with pyramidal cells (i.e., normal to the cortical meshes or using some more sophisticated estimation of the orientation of cortical columns (Bonaiuto et al., 2020)) and using volumetric sources estimations with free dipole orientation for subcortical regions (Attal and Schwartz, 2013).

EEG sources estimation in infants is relatively infrequent in the literature, and most studies that used it relied on volumetric approaches such as CDR (Xie and Richards, 2017) or ECD (Ortiz-Mantilla et al., 2019). It is more frequent in MEG infant studies (Kao and Zhang, 2019), but generally uses oversimplified spherical models to overcome the absence of realistic head models (Imada et al., 2006; Kuhl et al., 2014) or uses custom-built subject-specific realistic head models that are not reusable by the research community (Ramírez et al., 2017; Travis et al., 2011), forcing every research team to go through the time- and resource-consuming process of gathering infants MRI and building head models from scratch. The Neurodevelopmental MRI Database (NMD; (Richards et al., 2016) contains BEM and FEM realistic volumetric models of the head that can be used to perform source estimations using software that can use volumetric segmentation directly, such as FieldTrip (Oostenveld et al., 2011). However, resources such as FieldTrip^1^, Brainstorm (Tadel et al., 2011), and MNE-Python (Gramfort et al., 2013) which use surface models for cortical source reconstruction rely on external procedures for estimating the required cortical surfaces.

Reconstructing neuronal sources requires a few additional types of information to be combined with EEG recordings, including 1) a structural model of the head (conductor model); 2) a model of source distributions (source space); 3) the position of the EEG sensors on the subjects’ head (electrode placement); 4) a method for estimating scalp activity generated by neuronal sources (forward modeling); 5) an inversion scheme for estimating the probable neuronal sources corresponding to the observed scalp activity (inverse modeling). The first of these components, the structural head model, is generally built by post-processing magnetic resonance images (MRI) of the participants using specialized software designed for that purpose, such as FreeSurfer (Fischl, 2012), CIVET (MacDonald et al., 2000), the Computational Anatomy Toolbox (Gaser and Dahnke, 2016), BrainVISA (Rivière et al., 2009), SPM (Mattout et al., 2007), FSL (Jenkinson et al., 2012), or BrainSuite (Shattuck and Leahy, 2002). However, the time and expense associated with MRI scanning of the participants may be prohibitive or not possible with certain groups of participants because of health issues or ethical concerns. To address the lack of structural information in EEG analysis, population averages of head and brain structures have been proposed and can be used for computing approximate forward models (Fuchs et al., 2002; Valdés-Hernández et al., 2009). The objective of the current study is to extend these methods to infants by developing a novel set of 12 surface templates that can be used for source reconstruction for infants between 0 and 24 months of age.

## Method

### Volumetric dataset

To build age-specific templates that can be used for EEG source reconstruction using cortical surfaces, we used MRI averages available within the infants and preschool segments of the NMD version 2 (Richards et al., 2016), available on the NITRC website (https://www.nitrc.org/projects/neurodevdata). More specifically, we used the volumetric averages and four compartments boundary element method (BEM4) segmentation of head tissues contained therein.

The BEM4 volumes label every non-null voxel from head averages as belonging either to the skin, the skull, the cerebrospinal fluid, or the brain. In these segmentations, a minimum width of 1 mm was set for the skin compartment, 2 mm for the skull compartment, and 1 mm for cerebrospinal fluid compartment. These constraints help to avoid issues with intersecting surfaces when building source models. The identification of the brain, the skull, and the scalp on individual MRI was based on FSL Brain Extraction Tool (Bartlett and Smith, 1999; Smith, 2002).

Two sets of averages are provided, depending on whether MRI averaging was performed to optimize the head tissues or the brain. BEM classification is provided in the head averaging space. Table 1 lists the sample size per gender as well as the age range for the children included in every average. In this version of the database and for these age groups, all averages are based only on 3T scans using a T1 magnetization-prepared rapid gradient-echo (MPRAGE) sequence. See (Richards, 2013) for more details on individual MRI segmentation and (Sanchez et al., 2012) for volume averaging.

**Table 1.**
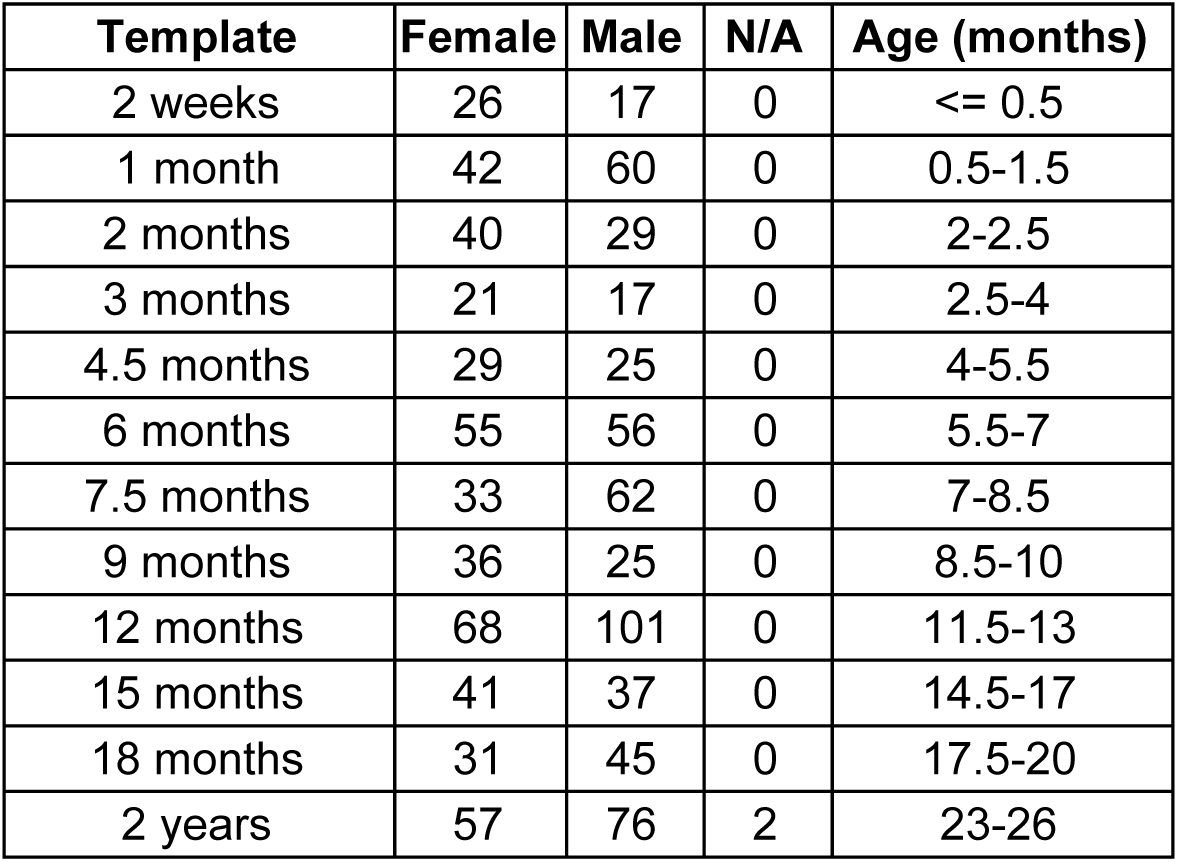
Sample size and ages for the different volumetric templates.

### Software

Aside from the extraction of brain surfaces relying on the Infant FreeSurfer pipeline (de Macedo Rodrigues et al., 2015; Zöllei et al., 2020), most of the data processing and analyses have been performed using custom code relying on various Python packages, the principal ones being connected-components-3d 1.5.0 (Kemnitz and Silversmith, 2020), Matplotlib 3.1.2 (Hunter, 2007), MNE-Python 0.20.0 (Gramfort et al., 2014, 2013), NiBabel 3.0.0 (Brett et al., 2019), Numpy 1.18.2 (Oliphant, 2006; Walt et al., 2011), Pandas 1.0.3 (McKinney, 2010; The pandas development team, 2020), Pillow 7.0.0 (Clark et al., 2020), PyCortex 1.2.0 (Gao et al., 2015), PyMeshFix 0.13.3 (Attene, 2010), Scikit-Image 0.16.2 (Walt et al., 2014), Scipy 1.4.1 (Virtanen et al., 2020), trimesh 3.5.12 (Dawson-Haggerty et al., 2020), and XArray 0.15.1 (Hoyer and Hamman, 2017).

### Surface extraction

We first processed the volumetric dataset to extract the brain and head surfaces needed for surface cortical source extraction. For brain surfaces, we normalized the intensity of the brain volumetric averages (FreeSurfer mri_nu_correct command) and converted them to a standard 256 × 256 × 256 space with 1-mm-side voxels (FreeSurfer mri_convert --conform command). Then, we used the Infant FreeSurfer reconstruction pipeline (infant_recon_all; de Macedo Rodrigues et al., 2015; Zöllei et al., 2020) to extract the surfaces separating the pia mater, the gray matter, and the white matter. During this process, the Desikan-Killiany (Desikan et al., 2006) and the Destrieux (Destrieux et al., 2010) cortical parcellations were also computed. We skipped the skull stripping step included in the Infant FreeSurfer pipeline since the brain averages from the NMD are already skull stipped.

For the head surfaces, we used the volumetric BEM4 classification from the NMD and ran a cube marching algorithm (scikit-image marching_cubes_lewiner function) followed by a Laplacian smoothing (trimesh filter_laplacian function) and a mesh decimation (MNE-Python decimate_surface function). Surface topological defects such as holes, inverted vertex normals, or vertices with fewer than three neighbors were corrected using custom Python code relying on external functions (MeshFix.repair from PyMeshFix; repair.fix_normals and Trimesh.remove_degenerate_faces from trimesh) and on code snippets borrowed and modified from various MNE-Python functions. Final meshes were checked for water tightness using trimesh. Further, one BEM volume had artifacts showing as small line segments over the background. We corrected these by zeroing any small separated cluster of non-null voxels using the connected_components function form connected-components-3d. Examples of the initial volumes and extracted surfaces using these two parallel pipelines are illustrated in Figure 1.

**Figure 1.**
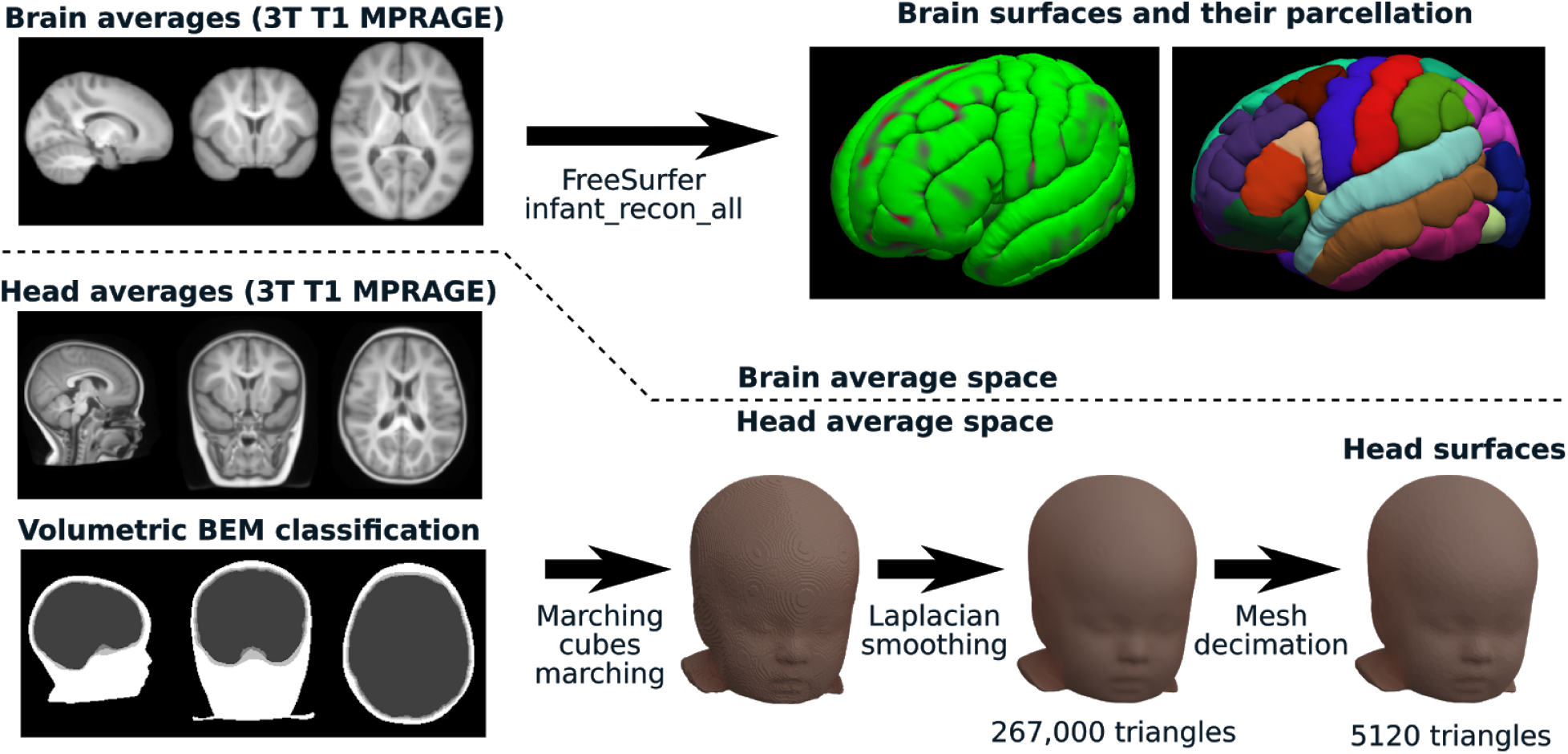
Surface extraction. On the left side, an example of the three types of volumes we used from NMD is shown. From this volumetric dataset, two parallel processes were run to extract the brain surfaces and their parcellation (top right; using Infant FreeSurfer) as well as the head BEM surfaces (bottom right; using custom Python code).

### Head-brain co-registration

Because the brain and head surfaces were computed from different volumetric averages, they were not initially perfectly aligned and needed to be co-registered. We aligned these surfaces manually using FreeView (a program from the FreeSurfer software suite) and custom Python code. An initial transformation (rotation, translation, scaling) was obtained by overlaying the head and the brain averages in FreeView (Figure 2.a). Further fine-tuning of translation and scaling was done in Python, using FreeView to visualize intermediate and final results (Figure 2.b).

**Figure 2.**
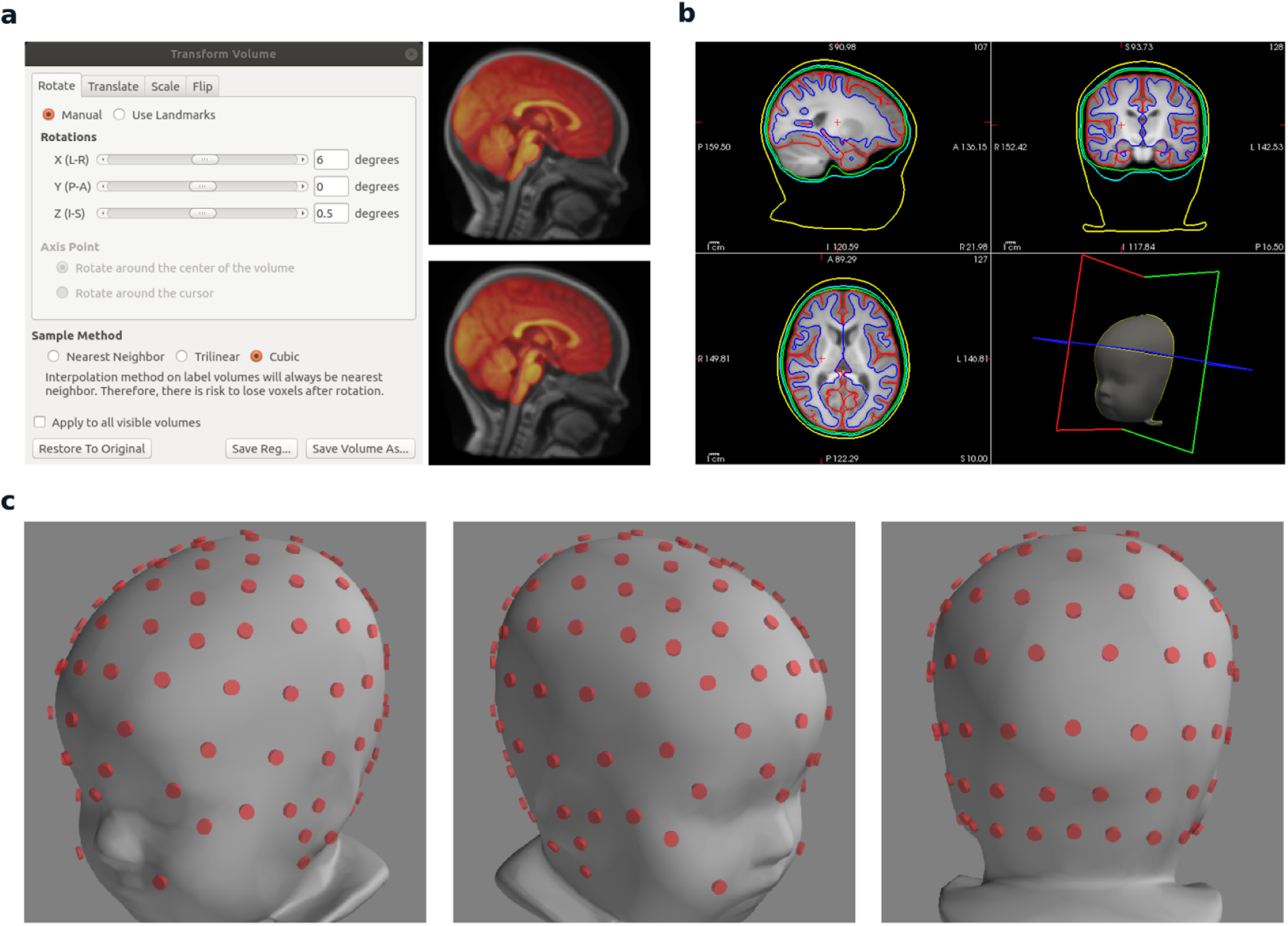
Co-registration processes. a) Initial head-brain co-registration by overlaying the brain over the head in FreeView, using partial transparency to help visual alignment. The brainstem offers very clear landmarks for finding the right rotation angle in the lateral view, as can be seen in the two examples of overlaid brains. b) Validation of alignment in FreeView after fine-tuning the scaling and the translation using Python code. For the forward model to work, the transform must be such that none of the surfaces (shown as color-coded lines: yellow: skin, cyan: outer skull, green: inner skull, red: pial surface, blue; grey-white matter border) intersect. c) Co-registration of sensors and head models using MNE-Python.

Once the transformation parameters were established (see Table 2), the transformation of the head surfaces to the brain space was done using:

**Table 2.** Head to brain space transformation parameters.

| | $t_x$ | $t_y$ | $t_z$ | $s_x$ | $s_y$ | $s_z$ | $\gamma$ | $\beta$ | $\alpha$ |
| --- | --- | --- | --- | --- | --- | --- | --- | --- | --- |
| **2 weeks** | 0.0 | -3.0 | 0.0 | 1.175 | 1.105 | 1.090 | -7.000 | -0.0 | 0.0 |
| **1 month** | 0.0 | -3.0 | 0.0 | 1.200 | 1.070 | 1.060 | -8.000 | -0.0 | 0.0 |
| **2 months** | 0.0 | -1.0 | 0.0 | 1.080 | 1.080 | 1.080 | -8.000 | -0.0 | 0.0 |
| **3 months** | 1.0 | 2.0 | 0.0 | 0.921 | 0.950 | 0.950 | 7.500 | -0.0 | 0.0 |
| **4.5 months** | 0.0 | 2.0 | -1.0 | 1.100 | 0.950 | 1.010 | 2.000 | -0.0 | 0.0 |
| **6 months** | -1.0 | -5.0 | -1.0 | 1.010 | 1.010 | 1.010 | 0.000 | -0.0 | 0.0 |
| **7.5 months** | -0.5 | -4.0 | 0.0 | 1.010 | 1.000 | 1.010 | -3.000 | 1.0 | 0.0 |
| **9 months** | -0.5 | -3.0 | -2.0 | 1.050 | 1.000 | 1.020 | -5.500 | -0.0 | 1.5 |
| **12 months** | 1.0 | -4.0 | 2.0 | 0.990 | 0.990 | 0.975 | -9.500 | -0.0 | 0.0 |
| **15 months** | 1.0 | -4.0 | 3.0 | 0.950 | 1.000 | 1.000 | -8.000 | -0.0 | 0.0 |
| **18 months** | 0.0 | 1.0 | 3.0 | 1.020 | 1.050 | 1.010 | 0.000 | -0.0 | 0.0 |
| **24 months** | 0.0 | 0.0 | 0.0 | 1.010 | 1.010 | 1.010 | -3.517 | -1.0 | 1.0 |

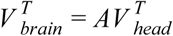

with *A* being an affine transform matrix (here rigid body plus scaling only), and *V* _*head*_ and *V* _*brain*_ being matrices of size N X 4, where N is the number of vertices in the mesh and each matrix row specifies the augmented vertice coordinates as [*x y z* 1].

The overall transform matrix is composed of elementary transforms as follow:

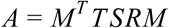

where M is the transform that map the coordinate system from neuroimaging RAS (right-anterior-superior) to the mesh vertices in LIA (left-inferior-anterior) coordinate system:

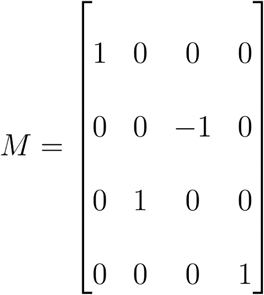

T is the translation matrix:

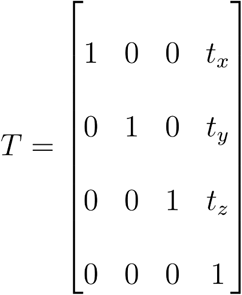

S is the scaling matrix:

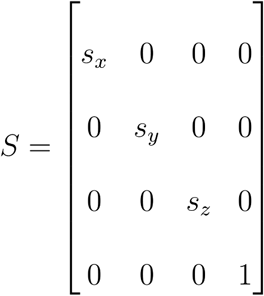

R is the rotation matrix, combining the yaw (*R*_*z*_(*α*)), the pitch (*R*_*y*_(*β*)), and the roll (*R*_*x*_(*γ*)):

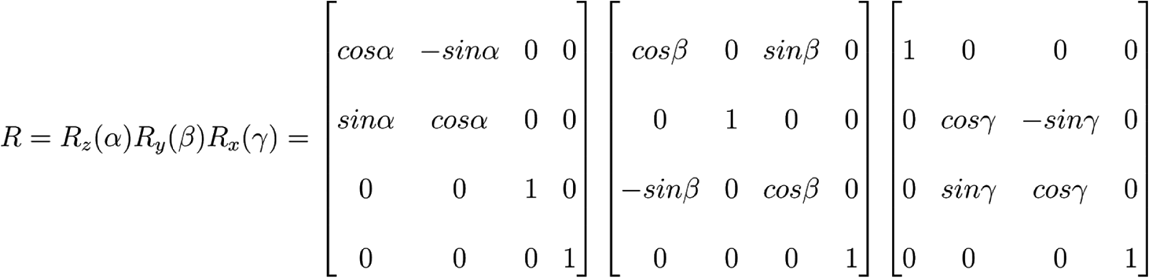

### Sensor co-registration

To map the scalp activity to neuronal sources, the EEG sensors must be placed over the scalp of the forward model. Electrode placement for the 3, 4.5, 6, 7.5, 9, 12, and 24 months time points were previously computed as described in (Richards et al., 2015) for the HydroCel GSN 128 channel sensor net and made available in the NMD. The electrode placement for the 2-weeks, 1- and 2-months templates were obtained from the 3-month average template electrode placement. This was done by registering the normative 10-10 positions from the 3-month-old average template to every participant of those age in the NMD, using the Coherent Point Drift method (Myronenko et al., 2006; Myronenko and Song, 2010), and translating the 3 months HydroCel GSN points into the participant space. Then the participant MRIs were registered to their age-appropriate average template and their electrode positions were appropriately translated into the average template space and averaged across participants. The averaged electrode positions were further scaled to fit to the head in the averaged MRI volume. The same procedure was used to fit the 12-months electrode placement to the 15- and 18-months time points. These placements being in the head average space, the head-to-brain transformation described in the previous section was applied to co-register them to the brain average space. The fitted electrode placement is provided with these templates for easy re-use. Other nets can be fitted following the same procedure.

### Validation dataset

To validate the use of the templates for source reconstruction, we used the 100 recordings (female=62; male=36; unknown=2) available in the 7-month-old/London segment of the International Infant EEG Data Integration Platform (EEG-IP) cohort (van Noordt et al., 2020). These EEGs were recorded using the HydroCel GSN 128 channel sensor net and a 500 Hz sampling frequency. They used an event-related paradigm with 1-second epochs starting at the moment when the visual stimulus was changed. The images used as stimuli were presented on a computer screen and consisted of a face looking directly at or away from the child, or a randomized noise control stimulus. For our analyses, we used the pre-processed recordings available in the EEG-IP cohort, in which automatic artifact rejections plus manual quality control has been performed using the Lossless pipeline (van Noordt et al., 2020).

### Source estimation

For comparing estimated sources, aside from our 12 infant templates, we also used the standard “fsaverage” from FreeSurfer, an adult template built using spherical surface averaging (Fischl et al., 1999) of the Buckner40 cohort which comprises 40 non-demented subjects (21 women) ranging in age from 18 to 30 and 65 to 93 years of age (FreeSurfer team, 2020a). Event-related EEG sources were computed using MNE-Python. The same parameters were used for source estimation using the 12 infant templates and the adult fsaverage template. We used the dSPM inverse operator and set the regularization parameters lambda2 to 1. Other parameters were left to their default value, as set by MNE-Python. The covariance matrix was estimated using the *method=“auto”* in the *mne*.*compute_covariance* function, which uses four different estimators (the Ledoit-Wolf estimator (Ledoit and Wolf, 2004) with cross validation for optimizing alpha, diagonal regularization, sample covariance, and factor analysis with low-rank (Barber, 2012)) and chooses the optimal solution base on log-likelihood estimation and cross-validation (Engemann and Gramfort, 2015).

### Data and code availability

The code for building the templates and for reproducing the validation analyses is available on GitHub (https://github.com/christian-oreilly/infant_template_paper). The volumetric averages and the final surface templates are available from the NMD (https://jerlab.sc.edu/projects/neurodevelopmental-mri-database/) and can be downloaded through the NeuroImaging Tools & Resources Collaboratory (https://www.nitrc.org/projects/neurodevdata). These templates are further being integrated to MNE-Python and Brainstorm for easier use.

## Results

### Templates and source reconstruction

We built surface-based structural templates at 12 time points between 2 weeks and 2 years of age (Figure 3). These templates have been tested using Brainstorm and MNE-Python and can successfully be used to reconstruct sources at these different ages (Figure 4).

**Figure 3.**
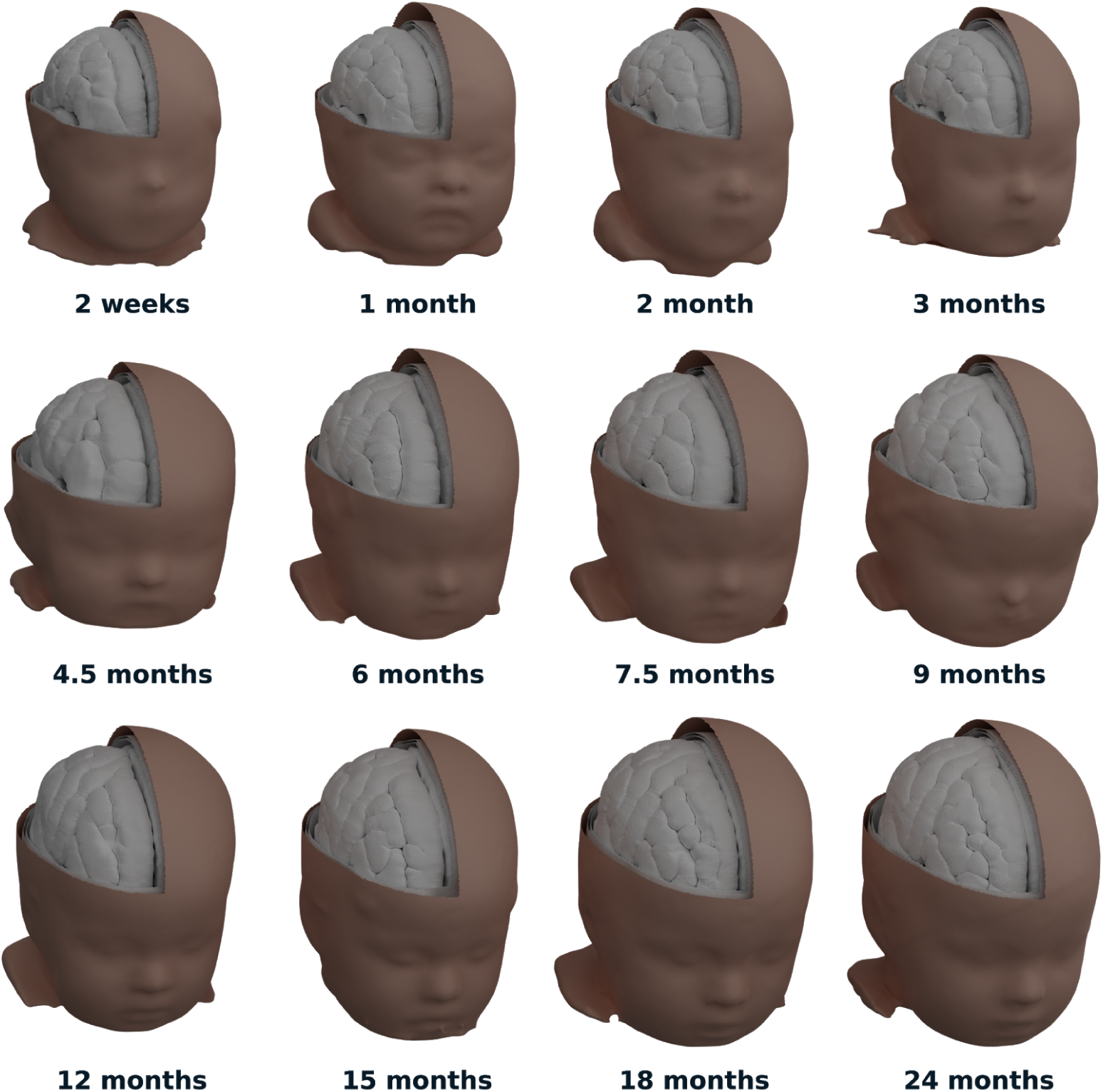
Twelve templates from 2 weeks to 2 years old comprising the surfaces for the scalp, the outer and inner surface of the skull, as well as surfaces separating the meninges, the grey matter, and the white matter (using non-decimated meshes and Blender rendering).

**Figure 4.**
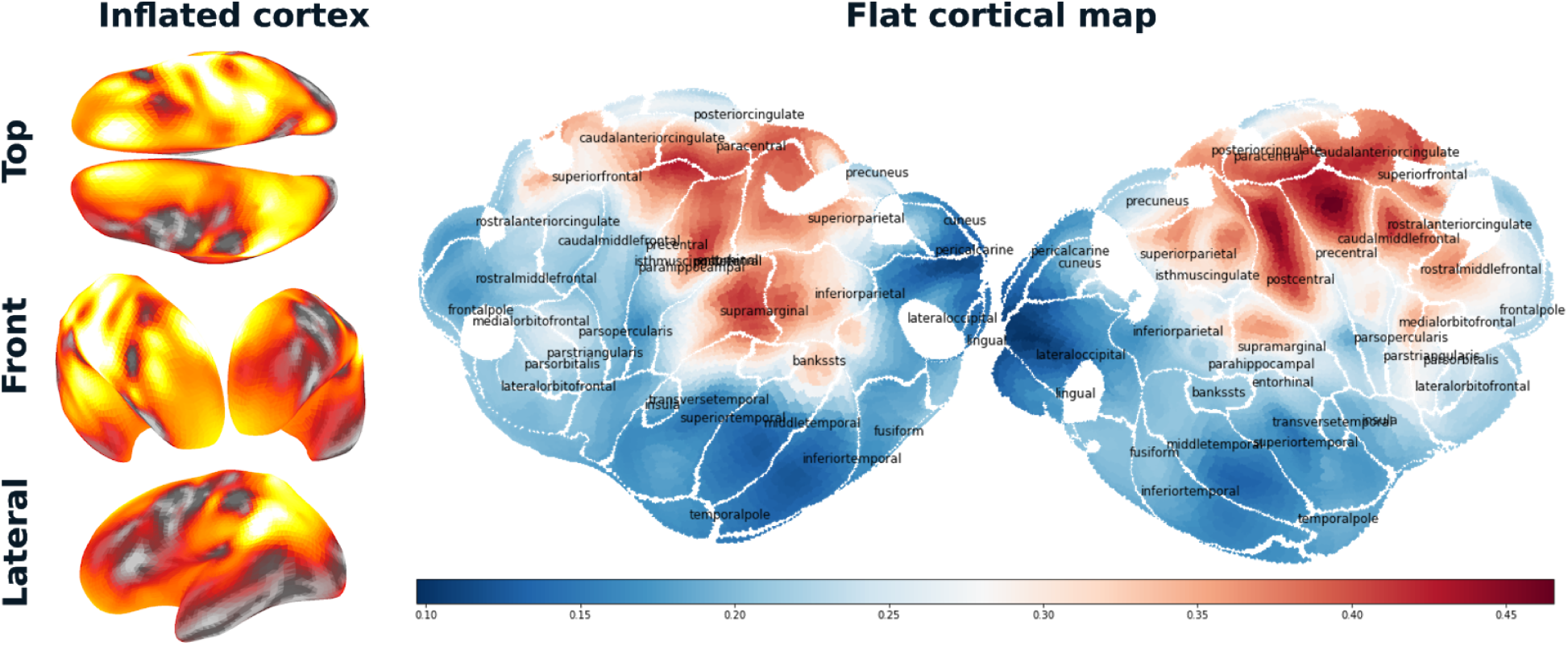
Example of source reconstruction using the 12-month template and showing the average source power over the 1-second epoch for the “noise” stimulus. The left panel shows inflated hemispheres, as provided by MNE-Python while the right panel shows the same source activity over the flattened cortical surfaces, using PyCortex.

### Validation

We validated the templates by testing the hypothesis that EEG sources should be more correlated when estimated from templates of similar age than from templates of very different ages. Thus, we computed correlations between the time-series of event-related EEG sources for pairs of templates and averaged across stimulus conditions, subjects, and brain regions. Heat maps in Figure 5.a,b show high average correlations between infant templates within the 0-2 years age range. However, there is a general tendency for values close to the diagonal to be smaller than those far from the diagonal, confirming that differences in estimated sources increases with template age differences. Further, the correlation with the sources computed using fsaverage is relatively low, confirming the large differences in source estimations when using an adult template for analyzing infants data.

**Figure 5.**
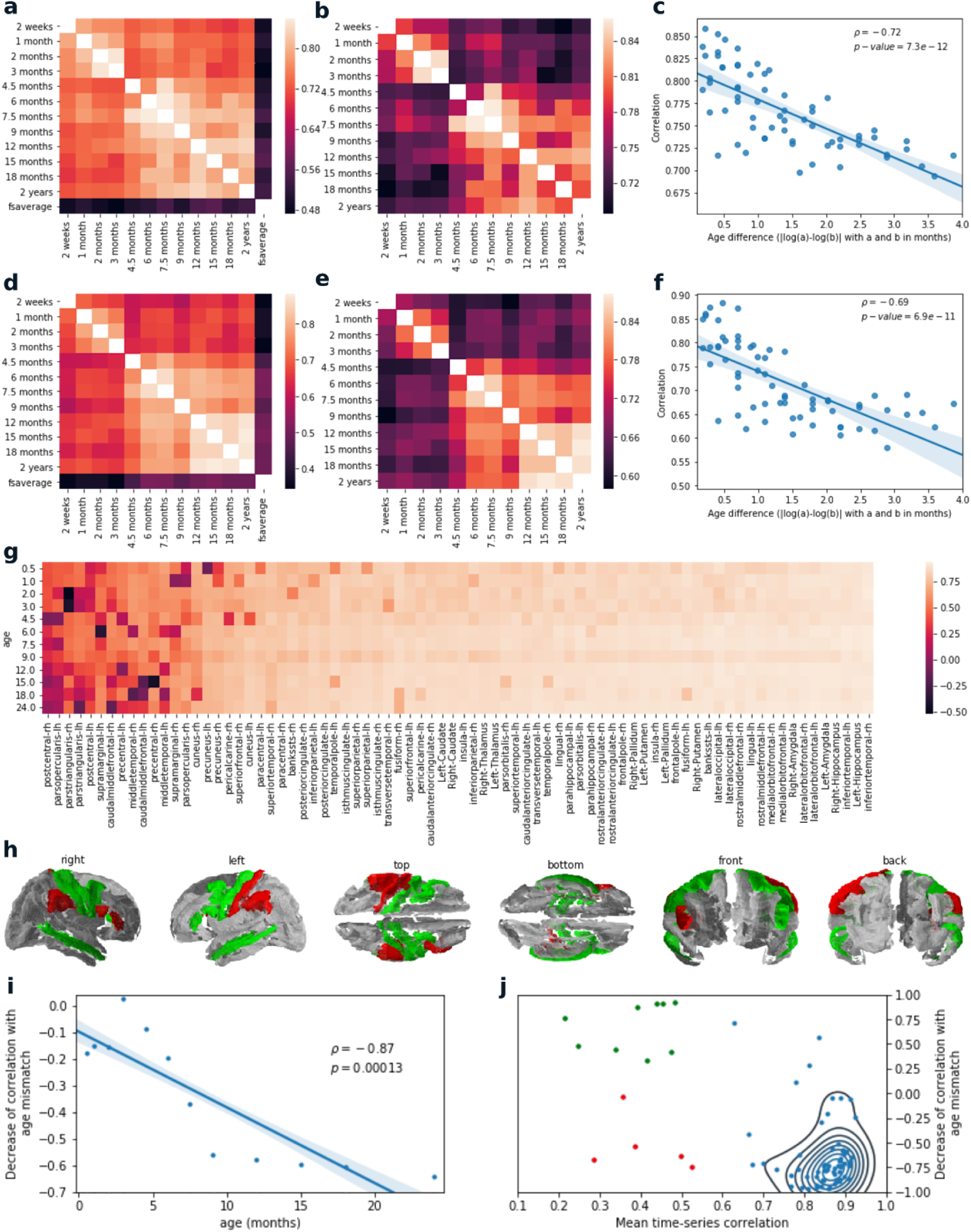
a) Average correlations between the source time-series computed from different templates, b) excluding the fsaverage template and adjusting the scale to emphasize the pattern of variation within the infant templates. c) Average time-series correlations as a function of the log-transformed age difference between templates. d-f) Similar as for panels a-c, but correlating across brain regions instead of time. g) Average time-series correlation computed separately for every region and averaged by reference template. h) Cortical representation showing average time-series correlation (mean values per column of panel g) lower than 0.6 colored in red or green, depending on whether they show or not the typical correlation decrease with increasing age differences. i) Value of the meta-correlation (i.e., correlations between age difference and the time-series correlations) with respect to the template age. j) Scatter plot of the average time-series correlation and meta-correlations for the different brain regions, color-coded as in panel h. All p-values are for one-tail tests.

The increasing difference between estimated sources with increasing template age difference is clearly demonstrated by linearly regressing the correlations plotted in Figure 5.a,b by the absolute differences between the logarithm of the template ages (R^2^=0.512, p-value=7.3e-12, N=66; Figure 5.c). Similar observations can be made by correlating sources across brain regions rather than across time-series (R^2^=0.477, p-value=6.9e-11, N=66; Figure 5, d-f). We further observed that the correlation of time-series between templates is generally very high across regions but is lower and more variable in some cases (Figure 5.g), particularly around the motor and somatosensory areas (Figure 5.h).

We refer to the correlations between the age difference and the time-series correlations as “meta-correlations”. They are computed by using the correlations associated with a given reference template (e.g., selecting a given column of the correlation matrix shown in Figure 5.b) and regressing these values against the age difference as shown in Figure 5.c. Such meta-correlations were computed across templates and regions (Figure 6.a) and then averaged across regions. Resulting averages were plotted against the age of the reference template (Figure 5.i). We observe meta-correlations with higher amplitudes as age increases, probably due to various factors complicating accurate modeling at younger age (e.g., less clear distinction between white and grey matter on infants MRI; normalization of non-fully gyrified brains using standards tools based on adult gyrification). The region and the template age together account for around 35% of the total variance of the meta-correlations (Table 6).

**Table 6.** Analysis of variance (ANOVA) for the linear model describing the variation of meta-correlations as a function of the age difference (continuous factor), the sample size (continuous factor), and the brain region (categorical factor). R^2^=0.35; adjusted R^2^=0.29.

|  | Sum sq. | df | F | PR(>F) |
| --- | --- | --- | --- | --- |
| region | 18.78 | 79 | 1.90 | 9.64E-06 |
| age | 38.57 | 1 | 308.46 | 2.40E-59 |
| Residual | 108.91 | 871 | — | — |

**Figure 6.**
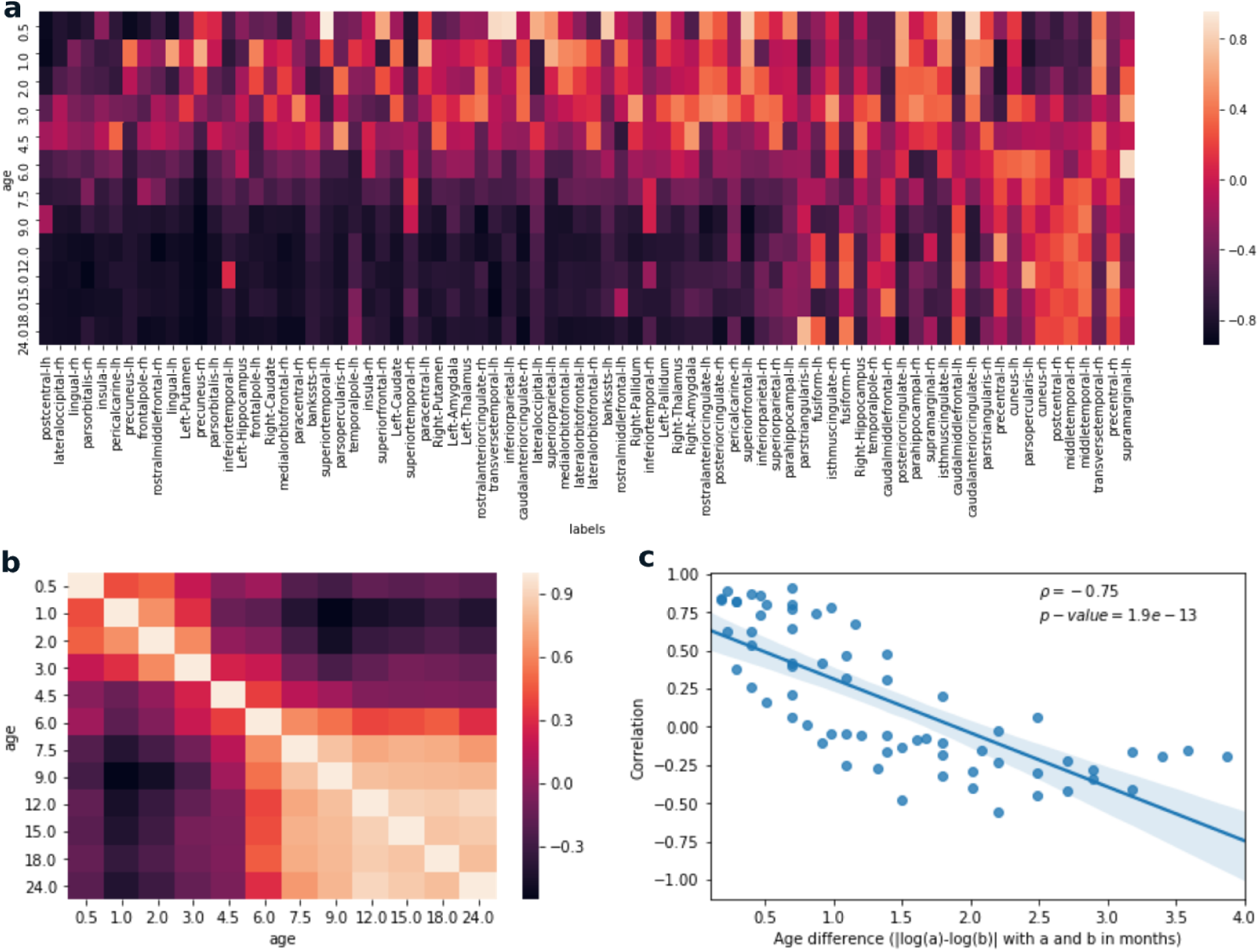
Meta-correlations correlating age differences with time-series correlations. a) Meta-correlations with respect to template ages and brain regions. b) Matrix of the correlation coefficients between rows of the matrix displayed in panel a. c) Values of the matrix from panel b plotted against age differences. Correlation coefficients and corresponding p-values (one-tail) are shown in the upper right corner of the graph.

For most regions, both the time-series correlations and corresponding meta-correlations with age differences are elevated but a small group of regions shows significantly lower (i.e., < 0.6) time-series correlations and a subgroup of these regions with lower correlations also shows meta-correlations with an inverse sign (Figure 5.j).

Figure 6.a shows that meta-correlations for various brain regions differ for the younger (up to about 5 months) and the older reference templates. The differences are even clearer when the correlation matrix between rows of the Figure 6.a are computed (Figure 6.b). Aside from showing a strong relationship with age difference (rho=-0.75; p-value=1.9e-13; Figure 6.c), it clearly shows the two aforementioned clusters (Figure 6.b) separating younger and older templates around 5 months of age.

## Discussion

A reliable estimation of EEG cortical sources requires the use of individualized or, when not available, population-averaged structural templates of the head. Standard automated approaches developed for building such templates from adult MRI scans do not provide satisfactory results in the pediatric population due to a lower contrast between white and grey matter (Phan et al., 2018; Schumann et al., 2010). Further, most automated pipelines rely on co-registering individual MRI against a population average, which may cause significant errors when using an adult average for co-registering younger populations. With some manual intervention to guide or correct white matter and grey matter classification, FreeSurfer can provide satisfactory results in children who are five years or older (Schumann et al., 2010) but not younger (FreeSurfer team, 2020b). For that reason, an Infant version of the FreeSurfer pipeline has recently been developed that covers the 0-2 years age range (Zöllei et al., 2020). To accelerate progress in EEG neurodevelopmental studies, we used this pipeline to develop 12 new templates for infants in this age range. These templates can be used for source reconstruction with software relying on surfaces of the brain and the head for their forward models. For approaches relying on volume models (e.g., to estimate volumetric current density) instead of surfaces, volume BEM and FEM models are already available in the NMD, which will allow comparing and benchmarking these different approaches for infants in the future.

The age difference within the 0-2 years range resulted in relatively mild differences in estimated EEG sources, suggesting that the reference templates are fairly accurate. Improvements in accuracy of source estimates may nevertheless be useful for some of the regions that showed lower and more variable correlations between templates (Figure 5.g). In the absence of quantitative resources that can be queried across age and brain regions for maturity index, it would be difficult to establish whether these lower correlations are due to faster maturational changes in these regions within the age range covered by these templates. We nevertheless note that most of these regions are around the motor and somatosensory areas, which are known to develop at a fast rate in the first few months following birth (Ganzetti et al., 2014), whereas the correlations are reliably high in frontal associative regions (Figure 5.h), which are known to develop later in life (Sowell et al., 1999). Some of these regions with lower correlations also show meta-correlation with an inverse sign (i.e., time-series correlation increasing with age mismatch; Figure 5.k), an observation that is unexpected and that is occurring either due to randomness or to an unknown cause.

In future, increased surface template accuracy may be achievable by using a surface-based registration directly rather than extracting surfaces from a volume-based average (Ghosh et al., 2010). This method might result in reference templates for infants with inflated cortexes and flat cortical maps closer to results obtained from adults. It is unclear, however, if such improvements would translate into a significant improvement regarding EEG source estimation since the gain obtained by increasing the template accuracy may be insignificant compared to the difference between the population average template versus the brain of each individual participant. It would also be interesting to investigate the extent to which the use of different templates would affect source imaging in studies using magnetoencephalography (MEG) instead of EEG, given that these two modalities measure fundamentally related but complementary components of the electro-magnetic fields produced by cerebral activity (Hämäläinen et al., 1993; Sharon et al., 2007).

Up to recently, no open-access population average surface models were available to perform cortical source reconstruction in infants. By releasing the 12 infant templates described in this paper, we are making source reconstruction based on cortical surfaces accessible for situations where individual MRI are not available and population averages are required, addressing a clear need in the EEG and MEG community. We are currently working at integrating these templates directly within MNE-Python and Brainstorm to allow researchers to easily benefit from them. This resource will undoubtedly come particularly handy for cross-sectional and longitudinal studies of typical and atypical neurodevelopment.

## Acknowledgments

This research is supported by the Azrieli Centre for Autism Research (ACAR) and NIH-R01NS104585 (EL; PI: M. Hamalainen). The computational infrastructure was provided by Calul Quebec (www.calculquebec.ca) and Compute Canada (www.computecanada.ca).

## Abbreviations

BEM: boundary element method
CDR: current density reconstruction
EEG: electroencephalography
EEG-IP: International Infant EEG Data Integration Platform
FEM: finite element method
MPRAGE: magnetization-prepared rapid gradient-echo
NMD: Neurodevelopmental MRI Database
MRI: magnetic resonance imaging

## Footnotes

1 FieldTrip can use surfaces for its source model, with the surfaces constructed from external tools such as iso2mesh (Fang and Boas, 2009), Brain2Mesh (Tran and Fang, 2017), or FreeSurfer (Fischl, 2012).

